# Melanin Nanoparticles as an Alternative to Natural Melanin in Retinal Pigment Epithelium Cells and Their Therapeutic Effects against Age-Related Macular Degeneration

**DOI:** 10.1101/2022.02.22.481315

**Authors:** Yong-Su Kwon, Min Zheng, Alice Yang Zhang, Zongchao Han

**Author notes:** Correspondence to: Zongchao Han.

## Abstract

Melanin is a natural pigment that is widely distributed in many parts of the human body, such as the skin and retinal pigment epithelium (RPE) in eyes. In contrast to skin melanin, which is being constantly synthesized by the epidermal melanocytes, melanin in the RPE does not regenerate. Melanin is known to function as a potential radical scavenger and photoprotective agent. However, the protective effects of melanin against oxidative stress decline with increasing age. This phenomenon has been significantly correlated with the pathogenesis of age-related macular degeneration (AMD). To increase the potential antioxidant and photoprotective characteristics of the melanin, we designed a new strategy by replenishment of melanin using PEGylated-synthetic melanin nanoparticles (MNPs) in the RPE for the treatment of AMD. We performed experiments using AMD-like cellular and mouse models and demonstrated that MNPs are safe, biocompatible, and selectively target reactive oxygen species (ROS) with powerful antioxidant properties. MNPs can traffic and accumulate in the RPE and are exclusively located in cytosol, but not the nucleus and mitochondria of the cells, for up to three months after a single-dose intravitreal (IVT) injection. Our findings demonstrate, for the first time, that MNPs are able to substitute for natural melanin in the RPE of eyes and suggest the potential efficacy of MNPs as a natural radical scavenger against oxidative stress in ROS-related diseases, such as AMD.

**Graphical Abstract:** 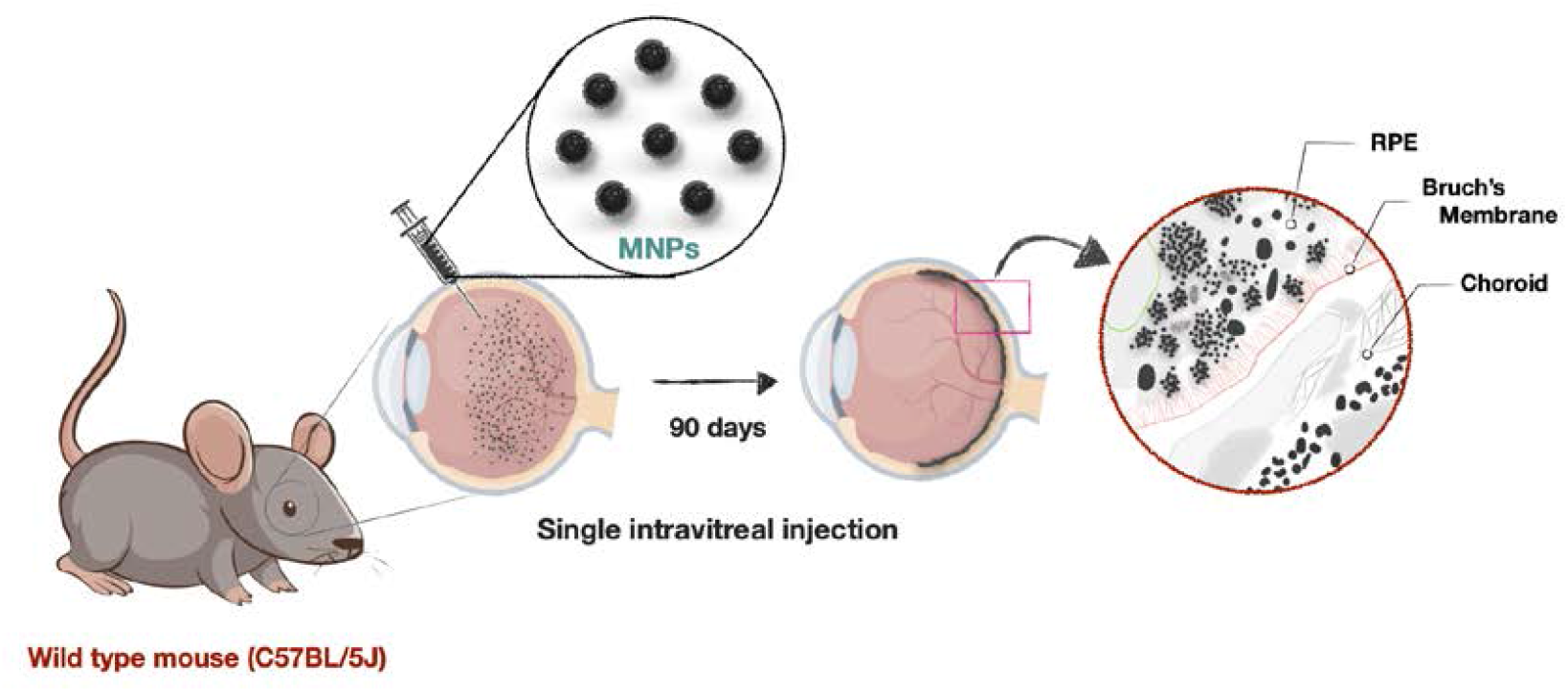

Intracellular trafficking showing that PEGylated-synthetic melanin nanoparticles (MNPs) reach their target and are stable in the retinal pigment epithelial cells for up to 3 months via a single-dose intravitreal (IVT) injection.

## 1. Introduction

Age-related macular degeneration (AMD) is a leading cause of blindness in elderly patients with a well-known public health impact, which is classified into two main pathological conditions: the non-exudative (i.e., dry) and exudative (i.e., wet or neovascular) forms.^1^ Despite significant advances in therapeutic options, the AMD incident rate has continued to grow.^2^ As current treatments for dry AMD are ineffective, there is a major need for therapies for dry AMD. Although treatments for wet AMD are effective, they are costly and are associated with a heavy burden of health care, and therefore there is a major need for improvements in treatments for wet AMD.^3^ The retina is one of the highest oxygen consuming tissues in the human body. The increasing production of reactive oxygen species (ROS) in the retina is affected by continual exposure to light, high concentrations of polyunsaturated fatty acids, and intensive oxygen metabolism.^4^ The overproduction of ROS by chronic oxidative stress can exceed the antioxidant capability of the retina and lead to damage of carbohydrates, membrane lipids, and proteins and nucleic acids in the retina, which are significant contributors to the development of AMD.^5, 6^ Increasing evidence shows that oxidative stress has played a decisive role in the progression of both wet and dry forms of AMD, representing a potential therapeutic target for AMD.^7, 8, 9, 10, 11^ Hence, a long-term sustained delivery of antioxidants for neutralizing oxidative stress may lead to a novel treatment targeted to preserve the retinal pigment epithelium (RPE) and photoreceptor cells in AMD.

Melanin is a well-known natural antioxidant, which plays a key role in protecting retinal cells from light-generated free radicals.^12^ Melanin is synthesized by L-DOPA, a precursor for the formation of melanin, during embryogenesis in RPE cells. However, it does not regenerate on adult RPE cells.^13, 14, 15^ The antioxidant activity of melanin in the RPE cells declines with advancing age.^16, 17^ Thus, maintaining the level of healthy melanin in RPE cells is very important because RPE cells are vulnerable to detrimental effects from aged melanin.

Very convincing evidence and clinically relevant data now support that nanomaterials have shown great promise as potential new tools in therapeutic approaches for ocular diseases.^18, 19^ Recently, dopamine-derived synthetic melanin nanoparticles have been suggested as a novel nanoantioxidant in biomedical applications with its highly biological relevance to natural melanin.^20, 21, 22^ We investigate the trafficking pathway and the final destination of PEGylated-synthetic melanin nanoparticles (MNPs) in biological systems and determine whether MNPs can be an alternative for natural melanin of RPE cells in vitro and in vivo. Our results provide strong evidence for their roles in attenuating pathological damages in AMD models. To the best of our knowledge, the melanin-based nanoantioxidants have not been studied in ophthalmic application yet, and there is still plenty of opportunity for discovery. We developed a safer and more effective antioxidant that can potentially achieve long-term effect through a single-dose administration to relieve pathological damages for AMD.

## 2. Results

### 2.1 Synthesis and characterization of synthetic melanin nanoparticles

The synthetic melanin nanoparticles (Bare-MNPs) were produced by oxidative polymerization of dopamine hydrochloride in the presence of sodium hydroxide as previously reported with slight modifications.^23^ The generation of Bare-MNPs were observed by the color of the solution that immediately turned to pale yellow and gradually became dark brown. Subsequently, thiol-terminated methoxy-polyethylene glycol (mPEG-SH) was introduced on the surface of Bare-MNPs by linkage via Michael addition reaction^24^ to improve the physiological stability of Bare-MNPs (Figure 1a). After the reaction, the prepared MNPs were observed to have an average diameter of ~ 80 nm with uniform nanospheres from transmission electron microscopy (TEM), which did not show any morphological difference after surface modification (Figure 1b, See Figure S1 in the supporting Information). Hydrodynamic size was measured by dynamic light scattering analysis (DLS) to determine the size distribution of nanoparticles. The DLS size of MNPs was ~ 100 nm and slightly increased to ~ 4.5 nm or less after surface modification compared to corresponding Bare-MNPs (Figure 1c). The zeta-potential analysis of these nanoparticles observed negative values, which was attributed to the multiple catechol groups on the melanin surface. Moreover, the resulting MNPs showed highly negative potential of - 48.2 mV (Figure 1d) due to the result of formation of a methoxy PEG layer with slightly negative charge on their surface, which was confirmed by Fourier transform infrared (FT-IR) spectra analysis (Figure 1e). The characteristic spectra peaks of MNPs showed the main absorption at 2880 cm^−1^ (the alkyl C—H stretching vibration) and 1110 cm^−1^ (the C—O—C stretching vibration) that were assigned to the PEG molecules. In terms of colloidal stability, MNPs revealed excellent dispersion ability in monkey vitreous for three months without particle aggregation (Figure 1f, Figure S2 supporting information) as well as exhibiting high colloidal stability in saline, cell culture media, and deionized water for a month (Figure S3 supporting information). These results validate that long-term stability of MNPs in biological media is suitable for biological applications.

**Figure 1.**
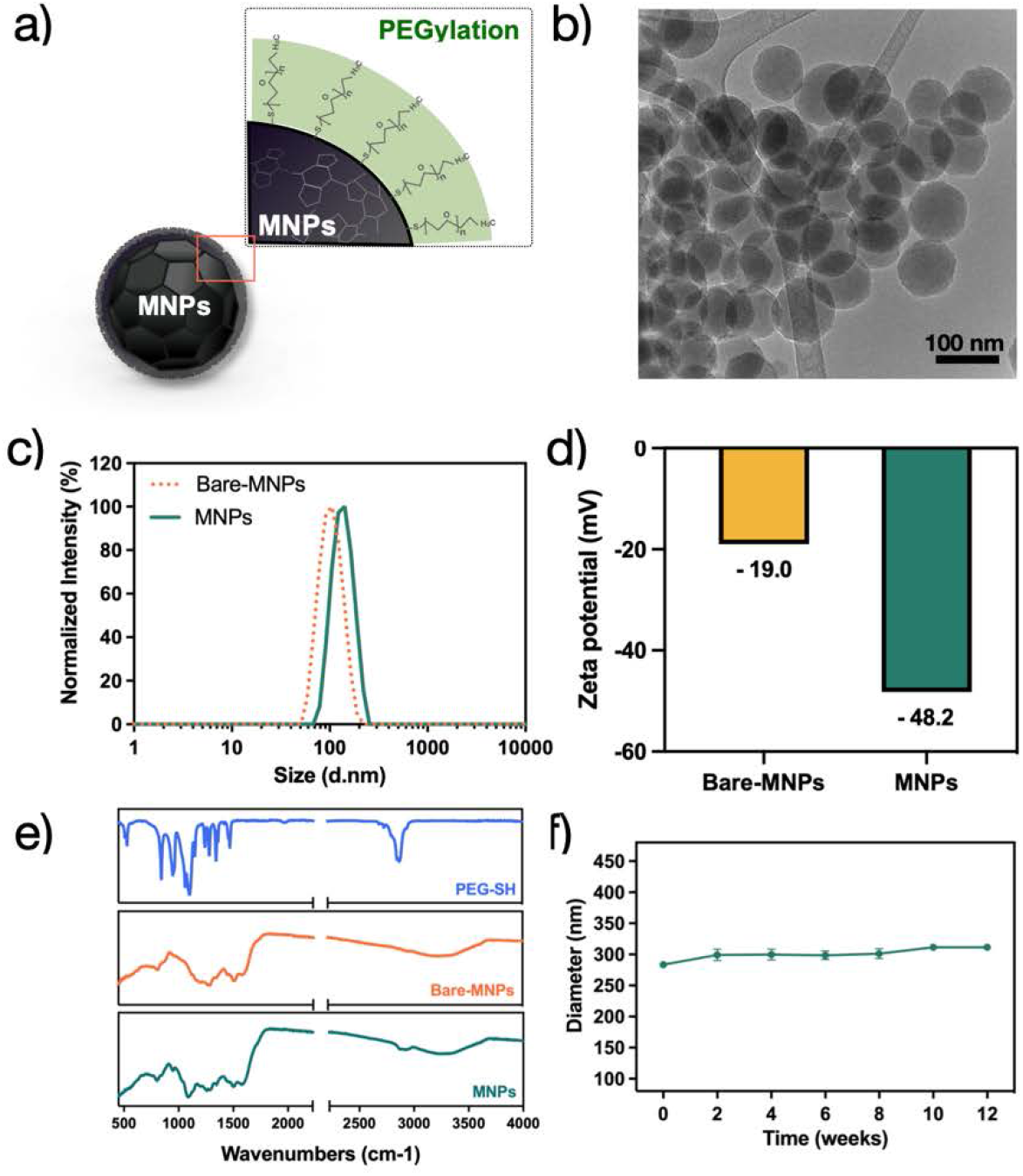
Characterization of nanoparticles. a) Schematic illustration of PEGylated-synthetic melanin nanoparticles (MNPs) structure. b) A transmission electron microscopy (TEM) image shows MNPs with spherical morphology and uniform size (~ 80 nm). c) Dynamic light scattering (DLS) analysis measures the hydrodynamic size of Bare-MNPs and MNPs to determine the size distribution of nanoparticles and relative monodispersity. d) Zeta potential of Bare-MNPs and MNPs. The results exhibited that the average surface charge was – 19.0 mV and – 48.2 mV, respectively. e) FT-IR spectra of mPEG-SH (2 KDa), Bare-MNPs, and MNPs. The characteristic spectral peaks of PEG appeared around 2880 cm^−1^ (alkyl C–H stretching) and 1110 cm^−1^ (C–0–C stretching). F) Colloidal stability of MNPs was measured in monkey vitreous for ~ 12 weeks.

### 2.2 Scavenging activity and intracellular uptake behavior of MNPs

Based on the distinct antioxidant properties of melanin in the biological system, we assessed the potential of MNPs to act as robust ROS scavengers. Here, we performed a 2, 2-diphenyl-1-picrylhydrazyl (DPPH) assay, which is widely used to predict the free radical scavenging capacity.^25^ DPPH produces purple color in methanol solution and fades to yellow color in the presence of antioxidants. We evaluated the capacity by monitoring the absorbance change at 516 nm. As shown in Figure 2a, MNPs exhibited excellent scavenging activity of ≈ 82 ± 2.5 % for free radicals at the dose level of 100 μg mL^−1^ compared with 0 μg mL^−1^ (PBS control). Moreover, the MNPs-mediated scavenging activity was verified through the concentration-dependent suppression behavior toward free radicals within our experimental dose range of 0 — 100 μg mL^−1^. We confirmed the highly sensitive antioxidant capacity of MNPs to scavenge free radicals. The result suggests that it shows free radical scavenging activity by their electron donating or accepting ability of MNPs.^26^

**Figure 2.**
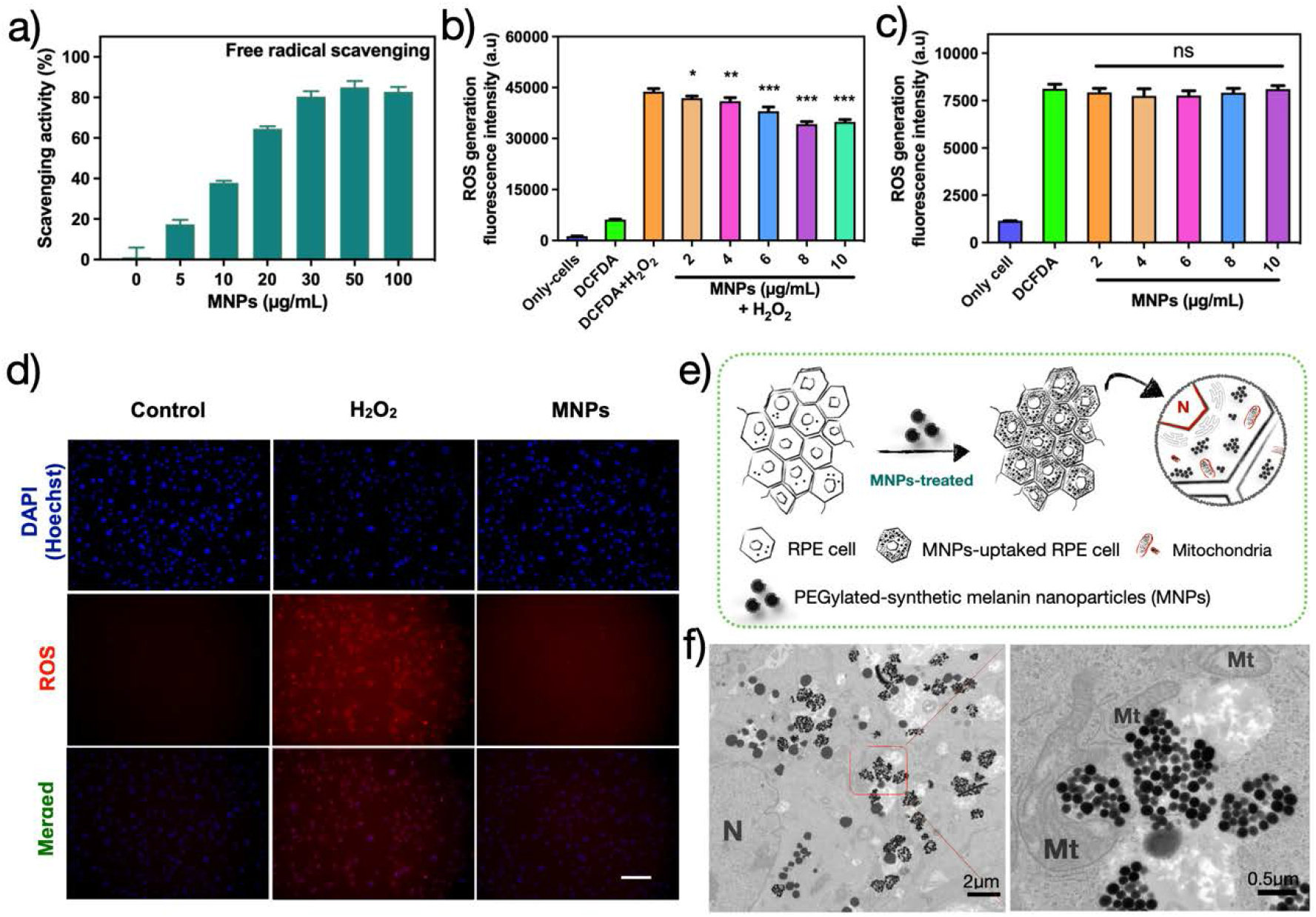
ROS-scavenging activity of MNPs in ARPE19 cells. a) The capacity of MNPs to scavenge free radicals by DPPH assay. b) Intracellular ROS scavenging activity of MNPs at different concentrations in H_2_O_2_-treated (0.575 mM) ARPE19 cells using DCFH-DA (50 μM). c) Intracellular ROS measurement following the same method as described above with no H_2_O_2_ treatment. d) Representative fluorescence microscopy images showing the ROS staining of ARPE19 cells under different treatments. Scale bar: 100 μm. e) Schematic illustration of MNPs trafficking and distribution in ARPE19 cells. f) Bio-TEM images show that MNPs present exclusively in cytosol of ARPE19 cells, but not in the nucleus and mitochondria (N; nuclear, Mt; mitochondria). ns, nonsignificant; *p <0.05, **P < 0.01, and ***P < 0.001 versus blank group, n = 4. All data are presented as the mean ± S.D.

### 2.3 MNPs-mediated protection on H_2_O_2_-induced oxidative stress cells

An imbalance between the production of ROS and the antioxidant defenses is implicated in persistent oxidative stress in RPE cells. Excessive accumulation of ROS leads to nonspecific inflammation and has been shown to be a potential factor in the pathogenesis of AMD.^6, 10, 11^ To determine scavenging ability of MNPs, we evaluated the amount of intracellular ROS levels in ARPE19 cells by DCF-DA fluorescence assay, which is used widely as a nonspecific indicator of ROS levels in cultured cells. We measured the intracellular ROS levels with 0.575 mM hydrogen peroxide (H_2_O_2_) that is sufficient to increase the significant amount of DCF fluorescence in ARPE 19 cells.^19^ H_2_O_2_-induced ROS production in ARPE19 cells was significantly suppressed by MNPs in comparison to untreated (0 μg) and DCF-DA (reagent only)-treated controls with the increase in the concentration of MNPs (Figure 2b). Intracellular ROS generation increased by about 7.5-fold upon addition of H_2_O_2_, whereas it clearly decreased upon treatment with MNPs (2 – 10 μg, *p <0.05, **P < 0.01, and ***P < 0.001). We further examined whether MNPs affected the fluorescence of the DCF-DA in absence of H_2_O_2_. The separated experiment showed that there were no interactions between DCF-DA and MNPs (2 – 10 μg, ns), which remained comparable to the fluorescence of DCF-DA (reagent only)-treated cells (Figure 2c). These results revealed that MNPs did not create any intracellular ROS. For all concentrations of MNPs, no obvious cytotoxicity was revealed under the experimental conditions, as observed through water-soluble tetrazolium salt (WST-1) assay (Figure S4 supporting Information). Additionally, ROS accumulation in the ARPE19 cells via administration of H_2_O_2_ (0.575 mM) were observed using MitoSox, a cell-permeable indicator that recognizes mitochondria-specific ROS. We confirmed that MNPs can be efficiently taken up into the ROS-induced ARPE19 cells (Figure S5 supporting information), and the exceeded ROS was significantly inhibited in the presence of MNPs (Figure 2d).

### 2.4 Visualization of MNPs in ARPE19 cells and ocular tissues

To evaluate whether MNPs reach their targets after administration, we performed TEM to visualize where the nanoparticles go both in vitro in ARPE19 cells and in vivo within the eye via intravitreal (IVT) injection (Figure 2e). The physiological levels of ROS in mitochondria play an essential role in the redox state and signaling in normal cellular metabolism. However, disruption of ROS balance in mitochondria can lead to the development of oxidative stress by excessive production of ROS that overflow to the cytosol from the mitochondria.^27^ To visualize the uptake pathway of MNPs by the ARPE19 cells, we used a Bio-TEM analysis with an epoxy resin embedding method. Compared with the un-treated group, we found that MNPs were exclusively located in cytosol after endocytic uptake and not in the nucleus and mitochondria of the transduced cells (Figure 2f), which were more prominent at various magnifications (Figure S6 supporting information). To investigate whether MNPs accumulation and distribution change in RPE, we injected 1 uL of MNPs (10 μg uL^−1^) via single-dose IVT injection and evaluated the trafficking of the nanoparticles at 3, 15, 30 and 90 days post-injection in wild type mice. Interestingly, we found that MNPs were exclusively accumulated in the RPE layer (Figure 3b), and their accumulation was gathering over time and they were not able to cross the Bruch’s (Br) membrane (Figure 3c-e). Similar to the in vitro trafficking study as described in Figure 2f, they were not found in the melanosomes and mitochondria in the RPE layer (Figure S7 supporting information), indicating that MNPs are capable of only targeting ROS that overflowed to the cytosol.

**Figure 3.**
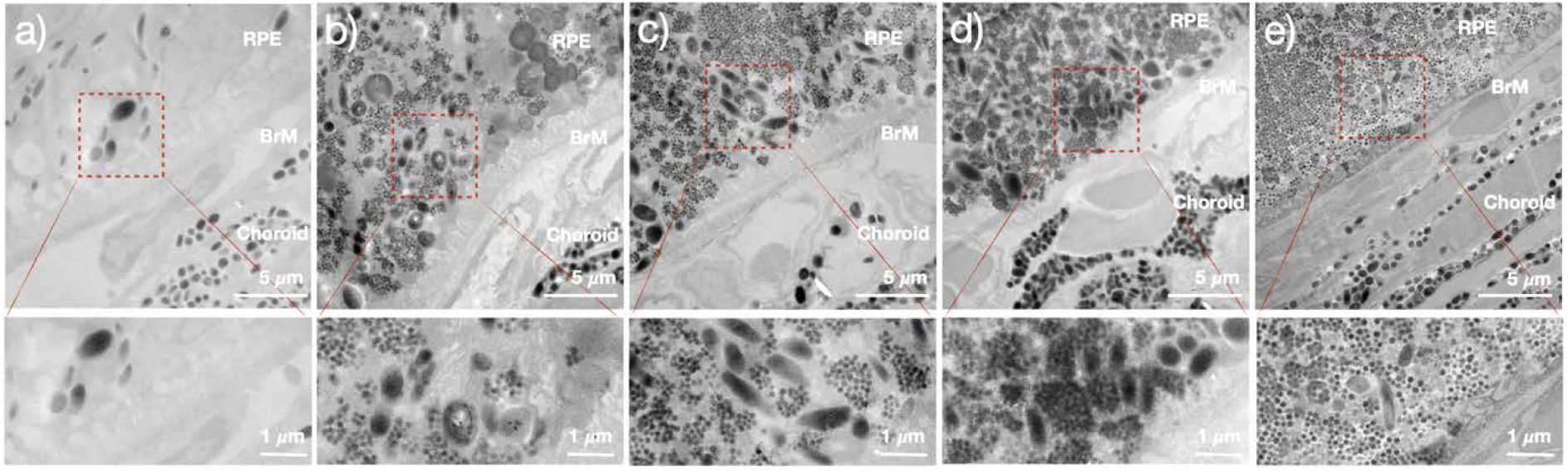
MNPs reach their target and stably remain in the RPE for up to 90 days via a single-dose intravitreal injection. Bio-TEM images show intracellular trafficking of MNPs. a) before and after IVT administration at b) 3 days, c) 15 days, d) 30 days, and e) 90 days, respectively. MNPs are clearly visible in the RPE layer. Inserts (bottom) are views at higher magnification as shown on the above. BrM: bruch’s membrane.

### 2.5 Neovascularization inhibition in laser-induced CNV mouse model

To demonstrate that the antioxidants of MNPs confer a benefit in AMD, we administered IVT injection of MNPs on a laser-induced CNV in C57BL/6J mice. The fundus fluorescein angiography (FFA) and optical coherence tomography (OCT) showed the presence of laser injuries immediately after the laser photocoagulation (top panels in Figure 4a, b and c). After FFA and OCT analysis of the CNV lesions (i.e., before injection), we performed an IVT injection with 1 uL of saline, aflibercept (40 μg μL^−1^), and MNPs (1 μg μL^−1^) to each laser-treated mouse eye, respectively. The FFA images showed that the MNPs-treated group clearly revealed a reduced vascular leakage from CNV lesions compared to the saline-treated group at 14 days after laser photocoagulation (bottom panels in Figure 4a and c). We also confirmed the presence of the rupture of Br membrane immediately after the laser photocoagulation and followed up retinal pathology with 2D cross-sectional OCT scans (indicated by arrow in Figure 4a, b and c). The OCT analyses showed that the thickness of laser-induced CNV lesions was significantly reduced in the MNPs-treated group compared to the saline-treated group at 14 days post-injection (Figure 4a and c). Similarily, the therapeutic effect of aflibercept (tradename EYLEA) was significantly higher than that of the saline-treated group. However, we observed that MNPs have more potent antiangiogenic activity than that of aflibercept, an FDA-approved drug which targets VEGF. A statistically significant difference was found between the two groups (p<0.01) (Figure 4b and c).

**Figure 4.**
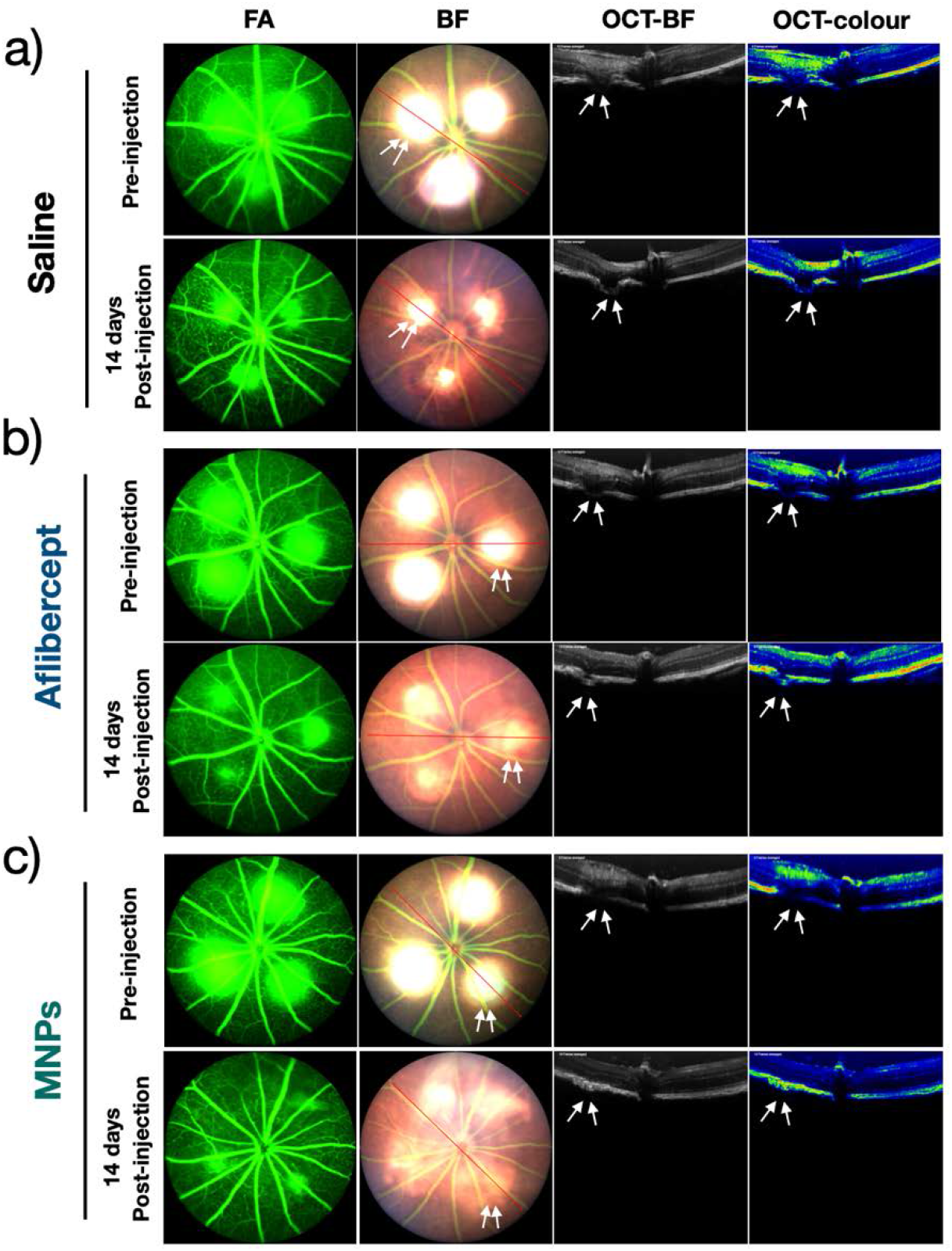
Fundus and OCT photograph analyses of choroidal neovascularization in a laser-induced CNV mouse model after IVT injection. a) Representative of fundus/OCT photographs in saline-treated eyes, which show just before injection (top panel) and after 14 days of laser injury (bottom panel). b) Representative of fundus/OCT photographs in aflibercept-treated eyes, which show just before injection (top panel) and after 14 days of laser injury (bottom panel). c) Representative of fundus/OCT photographs in MNPs-treated eyes, which show just before injection (top panel) and after 14 days of laser injury (bottom panel). White arrows indicate the areas of laser damage. Fluorescein angiography: FA, and bright field: BF.

The typical morphology of vascular area of laser-induced CNV lesions can be well quantitatively analyzed by choroidal flat mounts. To quantify the degree of CNV lesions, we prepared RPE/choroid/scleral flat mount specimens at 14 days post laser photocoagulation by staining with Alexa Fluor 488-conjugated isolectin-IB4 (IB4). IB4 is a commonly used endothelial cell specific marker. The areas of neovascularization were assessed using an Axiocam MR 5 camera on an Axio Observer.D1 inverted microscope (Carl Zeiss, Norway). We observed that the size of IB4-labeled CNV outgrowths was markedly reduced in the MNPs-treated group compared to that of the saline and aflibercept-treated groups (Figure 5a). The total area of CNV associated with each burn was measured per eye using Image J software (Figure 5b). The average size (in μm^2^) of CNV lesion per eye significantly decreased in the MNPs-treated group compared with saline- and aflibercept-treated groups (p<0.001). Thus, our results suggest that CNV lesions can be suppressed by antioxidant activity of MNPs.

**Figure 5.**
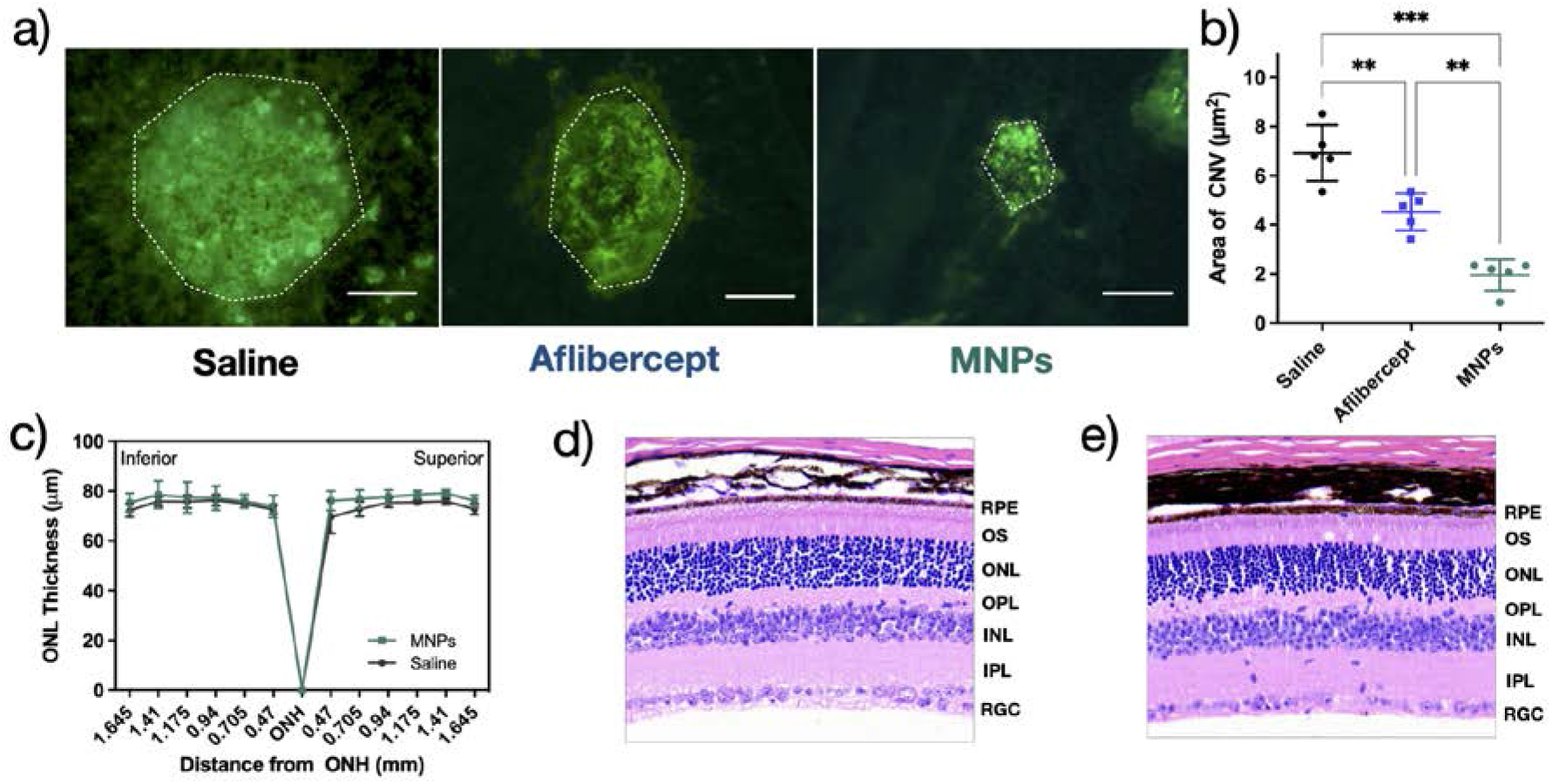
a) Representative images of RPE/choroid/scleral flat mounts from saline- and aflibercept-, and MNPs-treated eyes. Scale bar = 100 μm. b) Quantification of laser-induced CNV areas (μm^2^). Each dot represents one CNV lesion site. n = 5 lesions/group. Data are presented as mean ± SEM. one-way ANOVA followed by Tukey’s post hoc multiple comparison test, **P < 0.01, and ***P < 0.001. c) Quantification of outer nuclear layer (ONL) thickness. There are no significant changes in ONL thickness between saline- and MNPs-treated groups. Data are represented as mean ± S.D. Histological analysis of retinas in d) saline- and e) MNPs-treated groups.

### 2.6 Direct IVT delivery of MNPs is safe and has no toxicity to ocular tissues

To investigate whether IVT delivery of MNPs result in any toxicity or morphlogical changes in the eye, we conducted histologic analyses of the retinas. The outer nuclear layer (ONL) thickness serves as an important biomarker to monitor retinal pathological changes for visual function.^28^ Changes of ONL thickness are connected to toxicity to the ocular tissues. The measurement of ONL thickness in superior and inferior hemispheres was performed to evaluate the thickness changes after injection 1μL of saline and MNPs (1 μg μL^−1^) inside the vitreous space without laser photocoagulation. Mice were sacrificed at 5 days after injection and retinal cross-sections were prepared for histological analyses. Compared with saline-treated group, we did not find any morphological changes nor the effect on the retinal degeneration or any signs of inflammatory changes in the ocular tissues in the MNPs-treated group (Figure 5c, d, and e).

## 3. Discussion

In this study, we investigated the intracellular trafficking pathway and therapeutic capability of MNPs as a novel nanoantioxidant for AMD therapy. There is increasing evidence that suggests dysfunction of melanin contributes to the pathogenesis of AMD.^14, 15^ A study by Weiter et al. shows that there are about 2-fold higher density of melanin in the human macula area compared to that of the paramacular area in the RPE.^17^ It is well known that Caucasian people are more likely to develop AMD than African American people.^2^ This phenomenon may have been in part attributed by the different levels of melanin between the races. Studies have shown that compared with pigmented Abca4^-/-^ mice, early onset of oxidative stress and lysosomal impairment within the RPE were found in the albino Abca4^-/-^ mouse model of Stargardt macular dystrophy.^12, 29^ Indeed, a similar study was conducted in the mid-70s using congenic strains of rats which differ in melanin pigmentation.^30^ The post-mitotic RPE cells are derived from the neuroectoderm (neural tube), the embryonic precursor, which is known to develop into the central nervous system. The melanosomes in the RPE are synthesized during the prenatal period and show very little, if any, turnover after birth. An age-related decline occurs in either reduction of quantity or quality of melanin in the adult RPE. Conversely, lipofuscin (A2E), a major component of a toxic product known as “age pigment”, is found in AMD patients and is actively deposited on melanin granules throughout life.^31^ This phenomenon correlates with the stages of progression of lipid peroxidation and protein oxidation of cellular components in AMD pathology. Therefore, replenishment of melanin in the RPE in AMD patients could suppress retinal susceptibility to oxidative damage and represents a novel approach.

We prepared bioinspired Bare-MNPs using a simple coordination and self-assembly strategy, followed by PEGylation to create MNPs (approximately 80 nm) exhibiting excellent colloidal stability and biocompatibility with antioxidative activity against oxidative stress. Colloidal stability is a very important aspect in the evaluation for future experiments in clinical investigation. MNPs exhibited long-term colloidal stability in the monkey vitreous (for four months, last time checked) as well as the aqueous solution (for 12 months, data not shown) without particle aggregation. MNPs contain abundant hydroquinone residues that can serve as a trap for potentially harmful free radicals, which can protect cell damage from oxidative stress.^32^ We observed that MNPs can work as an efficient free radical scavenger as revealed by DPPH assay, which has a function similar to the eumelanin in the RPE, to protect ARPE19 cells against harmful oxidative stress without affecting cell viability.

Nanoparticles could easily pass through the biological barriers and enter the cells via pino- or macropinocytosis.^33^ We found that MNPs localize predominantly in the cytosol and do not enter the mitochondria or nucleus of transduced cells as described in Figure 2f. Given that mitochondria act as a source of ROS as redox signaling, we presume that it is much safer to access the intracellular ROS through scavenging in cytosol than that of directly going to mitochondria because it will not target the physiological levels of ROS in mitochondria that are essential for normal cellular metabolism but will only target the “bad ROS” that overflow to the cytosol. This phenomenon was further confirmed by *in vivo* trafficking study via a single-dose IVT injection in wild-type mouse. One of the most noticeable advantages of MNPs is that the nanoparticles prefer to accumulate in the RPE layer and do not pass through the Br membrane after a single-dose IVT injection for up to three months (last time checked). MNPs were evenly dispersed in the RPE layer similar to the *in vitro* trafficking studies, which do not enter nucleus or mitochondria in the RPE (Figure 2f, Figure S7 supporting information). The distribution behavior of the MNPs (i.e., remain in the cytosol and do not enter into the mitochondria or nucleus) is critical as this will allow cells to maintain their normal cellular metabolism and to keep their nucleus genetic material integrity without being influenced by the nanoparticles. These findings suggest that MNPs could function as a key factor to modulate the activity of melanin in the RPE without inducing any toxicity, providing insights into future directions using MNPs as a novel approach to overcome dysfunction of melanin. After confirming the trafficking pathway of MNPs in ocular tissues, we investigated the therapeutic effect of MNPs in a laser-induced CNV mouse model. Our results showed that MNPs could reduce the pathological injuries in CNV and are even superior to aflibercept, an FDA approved prescription medication, in the laser-induced CNV mouse model, indicating that MNPs play an important role as a robust antioxidant against harmful oxidative stress (Figure 4). Compared to major anti-VEGF agents (aflibercept, ranibizumab, and bevacizumab), that may be subject to issues of short half-life (7-10 days) and the effects of antibody uptake (e.g., diffusivity, antigen density, permeability, and systemic clearance).^34, 35^ The fact that the MNPs-treated retinas are structurally similar to that of wild-type control, as indicated by the histological analysis (Figure 5d, e), makes it as a superior antioxidant and a therapeutic tool to relieve pathological damage for AMD.

## 4. Conclusion

In conclusion, we demonstrated that MNPs can be used as a natural antioxidant defense platform for AMD therapy. The biocompatible MNPs exclusively accumulate in the RPE cells and exert excellent antioxidative effects, which help to alleviate pathological damages in AMD-like models. Our study takes advantage of the inherent antioxidant properties of melanin to combat oxidative stress and may lead to a novel treatment to preserve the RPE and photoreceptors in AMD.

## 5. Experiment Section

### 5.1 Synthesis of synthetic melanin nanoparticles

Bare-MNPs were synthesized according to a previously published method with minor modifications.^23^ In brief, dopamine hydrochloride (180 mg) was dissolved in 90 mL deionized water and then 780 μL NaOH (1N) solution was added, followed by gentle stirring at 50 °C. The mixture color instantly turned to pale yellow and gradually changed to dark brown. After reacting for 5 h, MNPs were purified with deionized water three times by centrifugation (Thermo Scientific™ Sorvall™ ST 16R centrifuge, USA) at 13,000 rpm for 15 min. The Bare-MNPs were finally redispersed in deionized water (5 mL) and stored at room temperature.

### 5.2 Surface modification of Bare-MNPs

10 mL of Bare-MNPs solution (1 mg mL^−1^) were adjusted to be pH 9-10 by adding NH4OH solution (28 wt%, Sigma-Aldrich) and then 50 mg of mPEG-SH (average Mn 2,000, Sigma-Aldrich) was added into the mixed solution under continuous stirring for 12 h. After the reaction was completed, the MNPs were collected by centrifugation at 13,000 rpm for 15 min, washing the pellet with deionized water three times. Characterization of MNPs was carried out by TEM, DLS, zeta-potential and FT-IR analysis as described in the Supporting Information 1.2.

### 5.3 Colloidal stability of MNPs

MNPs were prepared by centrifuging (13,000 rpm, 20 °C, 15 min) and redispersing the pellet in different solutions including monkey vitreous, cell cultured media, saline, and deionized water. The colloidal stability of MNPs was measured by DLS analysis at intervals of a week.

### 5.4 Free *radical scavenging assay*

To measure free radical-scavenging capacity of MNPs, DPPH radical scavenging assay was carried out with slight modifications. We prepared a fresh stock solution of ethanol containing DPPH (0.1 mM) solution and added a certain amount of MNPs with the final concentration at 0, 5, 10, 20, 30, 50, and 100 ug mL^−1^. All solutions were placed in a dark condition for 20 min and the absorbance was monitored at 516 nm by a spectrophotometer (Molecular Devices Spectramax M5) every 5 min for 30 min. DPPH solutions at the same concentration containing no tested samples were used as controls. The DPPH scavenging effect was calculated by following equation:

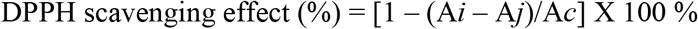

Where A*c* is the absorbance of control (without tested samples), A*i* is the absorbance of samples (with MNPs), and A*j* is the absorbance of the samples without DPPH solution (only MNPs). Samples were analyzed in triplicate.

### 5.5 Cell culture

ARPE19 cell line was used for in vitro studies. The cells were cultured in T-flask containing DMEM/F-12+GlutaMAX-I (Gibco, Cat. No. 10565018) supplemented with 10 % fetal bovine serum (FBS, Sigma, and Cat. No. 151400122) and 1 % antibiotic (penicillin/streptomycin) solution and incubated in a humidified incubator (Thermo Fisher Scientific, USA) at 37 °C under 5 % CO_2_ atmosphere.

### 5.6 Cytotoxicity of MNPs in ARPE19 cells

A WST-1 assay was carried out to measure cell viability, following the manufacturer’s protocol. For the experiment, ARPE19 cells were seeded in 96 well plates at a density of 8 X 10^3^ cells per well (n=4) and allowed to grow at a humidified cell culture incubator for 24 h. The cells were then exposed to 200 μL of fresh media containing different concentrations (0, 5, 10, 20, 30, 50 and 100 μg mL^−1^) of MNPs. The plates were continuously incubated for different time periods (1 – 2 days). At the end of the incubation period, the WST-1 reagent (Abcame, ab65473, USA) was added to each well and additionally incubated at 37 °C for 2 h. Absorbance of cell culture medium was measured using a spectrophotometer at a wavelength of 450 nm with a background subtraction at 630 nm. The absorbance values of control (untreated cells) were used as a reference value for determining the cell viability (%).

### 5.7 Intracellular ROS measurement

Based on the demonstrated effectiveness of MNPs as robust ROS scavengers, the intracellular ROS level was detected by using Reactive Oxygen Species Assay Kit (DCFH-DA, Invitrogen™). This method uses a fluorogenic probe as a sensitive marker of the cellular oxidation process and indirectly determines intracellular ROS level in cells. Once the DCFH-DA penetrates through the cell membrane, it is readily deacetylated to DCFH, which is further oxidized by intracellular ROS to a highly fluorescent compound (fluorescent 2’,7’–dichlorofluorescein (DCF)) in the cells. ARPE19 cells were seeded into 96-well plates at a density of 8 X 10^3^ cells per well (n=4) for 24 h, followed by the treatment of MNPs in different concentrations (0, 2, 4, 6, 8, and 10 μg mL^−1^). After 24 h of incubation with increasing concentration, the cells were washed with serum-free media and loaded 100 μL fresh media containing 50 μM DCFH-DA solution to each well with remaining untreated cells as control. The plate was then incubated at 37 °C for 60 min. The cells were washed with 1 X DPBS and treated with 100 μL fresh media containing H_2_O_2_ (final concentration: 0.575 mM) to each well, and then incubated at 37 °C for 2 h. Only DCFH-DA treated and untreated cells were used as controls. The DCF fluorescent intensity was measured by a spectrophotometer (Molecular Devices Spectramax M5) with excitation and emission wavelengths at 485 and 530 nm, respectively. To reliably measure levels of ROS in live ARPE19 cells, 5 μM (final concentration) of a fluorogenic probe (MitoSoX, Thermo Fisher Scientific Invitrogen™, USA) was added to each well after MNPs incubation and H_2_O_2_ treatment. The cells were then washed twice with 1 X DPBS and measured intracellular ROS level in the live ARPE19 cells with excitation and emission at 510 nm and 580 nm, respectively.

### 5.8 Embedding and thin sectioning with ultramicrotome

To explore a behavior of MNPs in ARPE19 cells, the cells were incubated in 6 well plates at a density of 2 X 10^5^ cells per well and exposed to 2 mL of fresh media containing of MNPs (100 μg mL^−1^). After 24 h, ARPE19 Cells (with non-treatment and/or MNPs-treatment) were fixed in 2.5% glutaraldehyde (25%, Sigma-Aldrich) in sodium phosphate buffer (0.15M, pH 7.4) for 1 h at room temperature and stored at 4°C until next processed. After rinsing three times with sodium phosphate buffer (0.15M, pH 7.4), the specimens were post-fixed in 1% osmium tetroxide (microscopy grade, Sigma-Aldrich) for 1 h at room temperature. After washing with sodium phosphate buffer (0.15M, pH 7.4), the specimens were dehydrated (30%, 50%, 75%, 100%, 100%, 10 minutes each) with increasing concentrations of ethanol and embedded in Polybed 812 epoxy resin (Polysciences, Inc.) to make plastic blocks. The small specimen blocks were sectioned to 100 nm thickness by a diamond knife using a Cryo-ultramicrotome (Leica EM UC7, CHANL-UNC). The ultrathin sections were then collected on 200 mesh copper grids and stained with 4 % aqueous uranyl acetate (98% EM grade, Polysciences, Inc.) for 5 min. The visualization was aided using a Thermo Scientific™ Talos F200X transmission electron microscope operating at 200kV, and digital images were acquired using a Ceta camera on Velox-acquisition software.

### 5.9 Animals

We used postnatal age 6 - 8 weeks C57BL/6J mice (Jackson Laboratory, ME, USA) for all in vivo experiments. All animal treatments or procedures were carried out and maintained in accordance with National Institutes of Health animal care guidelines as approved by the UNC Institutional Animal Care and Use Committee (IACUC).

### 5.10 Laser-induced CNV mouse model

To obtain a laser-induced CNV mouse model, we followed the standard protocol. In brief, adult 6-8 weeks C57BL/6 mice (20 – 25 g) were randomly divided into the three groups; saline-treated, aflibercept-treated, and MNPs-treated groups (20 mice per group) and the animals were anesthetized by intraperitoneal administration of a mixture of ketamine (85 mg kg^−1^) and xylazine (14 mg kg^−1^) (Butler Schein Animal Health, Dublin OH, USA). 1% tropicamide (Bausch & Lomb Inc., Tampa, FL, USA) was then dropped topically into the pupils for dilation, and GenTeal lubricant gel (Alcon) covered the eye to avoid any eye dehydration and loss of ocular transparency. Three lesions around the optic nerve were then implemented using laser photocoagulation (532 nm, 380 mW, 80 ms), and the injury on the Br membrane was confirmed by the appearance of an air bubble sign using a Micron III retinal image-guided laser system (Phoenix Research Laboratories, CA, USA). To observe the detailed morphological changes in mouse eyes, optic coherence tomography (OCT) and fundus imaging tests were carried out as described in the Supporting Information 1.3. The IVT injections were carried out after two days of photocoagulation following earlier protocols. Briefly, a hole was first made on the sclera of each eye using a sterile 30-gauge (G) needle. 1 μL of MNPs (1 μg μL^−1^), aflibercept (40 μg μL^−1^), and saline solution was slowly injected into the vitreous cavity using a 35 gauge needle attached to a 10 μL nanofil syringe through the punctured hole of the scleral surface, and the position of the needles within the eye was visualized using a microscope (Carl Zeiss Surgical, Incorporated, Thornwood, NY, USA). In case of in vivo trafficking study, 1 μL of MNPs (10 μg μL^−1^) and saline solution were slowly injected into the vitreous cavity of wild type mice (Non-CNV mouse model), RPE/Choroid/Scleral tissues at 3, 15, 30, and 90 days post-injection were embedded following procedures as described above. The needle was gently removed from the eye and triple antibiotic ointment (Equate, Wal-Mart, Bentonville, AR, USA) was dropped on the eye surface to avoid any infections. After all surgeries, the mice were kept warm for 20 min and sacrificed to collect tissues at the end point of our study.

### 5.11 Flat mounts and histology scan

After performing the last OCT and FFA imaging, the mice were euthanized for preparing RPE/choroid/scleral flat mounts as previously described.^18, 19^ Briefly, the eyes of the mice were enucleated and immediately fixed with 4 % paraformaldehyde for 1 h at room temperature. After washing three times with PBS, the RPE/choroid/scleral tissues were isolated and stained with Alexa Fluor 488-conjugated isolectin-IB4 (10 μg mL^−1^, Invitrogen™) overnight. The IB4-labeled tissues were fully washed with PBS three times for 15 min and then prepared for 4 or 5 radial incisions. The tissues were flattened and mounted with the vitreous side up onto the slide. The images were visualized by Axiocam MR 5 camera on an Axio Observer.D1 inverted microscope (Carl Zeiss, Norway). For histologic analysis, the collected eyes were fixed with 4 % paraformaldehyde and embedded in paraffin, followed by sectioning and H&E staining at the QuantCenter 3DHISTECH, USA.

### 5.12 Statistics

All studies were triplicated for each group and the results were reported as the mean ± standard deviation (S.D.). GraphPad Prism 9.0 software (La Jolla, CA, USA) was used to analyze all data also for statistical and kinetic analysis. Multiple group comparisons were examined by one-way analysis of variance (ANOVA) to determine the statistical differences; differences were considered as statistically significant at a p-value lower than 0.05 (P < 0.05).

## Acknowledgements

The authors thank Cassandra Janowski Barnhart (UNC Institute for Global Health & Infectious Diseases) and Ahra Cho for their critical reading of the manuscript. The authors thank Amar Kumbhar (The Chapel Hill Analytical and Nanofabrication Laboratory, CHANL, UNC-CH) and Victoria J Madden (The Microscopy Services Laboratory, MSL, UNC-CH) for their kind help with the transmission electric microscopy imaging. This work was supported in part by the Bright Focus Foundation (M2019063, Z.H.) and the Edward N. & Della L. Thome Memorial Foundation (138289, Z.H.).

