## Supplementary material for "Melanin Nanoparticles as an Alternative to Natural Melanin in Retinal Pigment Epithelium Cells and Their Therapeutic Effects against Age-Related Macular Degeneration": MNPs_SI_Final.docx

**1. Experimental Section**

**1.1 Materials**

Dopamine hydrochloride (Sigma-Aldrich, H8502), Sodium hydroxide (NaOH, Sigma-Aldruch, S8045), Hydrogen peroxide solution (H_2_O_2_, Sigma-Aldrich, H1009), Poly(ethylene glycol) methyl ether thiol (mPEG-SH, Sigma-Aldrich, 729140), 2,2-Dipheny-1-picrylhydrazyl (DPPH, Sigma-Aldrich, D9132). All chemicals were used as received without any further purification. Deionized water was used in all experiments.

**1.2 Material Characterization**

Transmission electron microscopy (TEM) (Thermo Scientific^TM^ Talos F200X) was used for MNPs morphological characterization at an accelerating voltage of 200 kV at the Chapel Hill Analytical and Nanofabrication Laboratory (CHANL, UNC-CH) and the Microscopy Services Laboratory (MSL, UNC-CH). TEM images were recorded after samples were collected on the 200-mesh carbon-coated copper TEM grids. To support TEM data and explain the size change of the nanoparticles, particle size distribution and zeta potential were analyzed by dynamic light scattering (DLS) using a Nano-ZS Zeta Sizer system (Malvern, USA). PEG functionalization on the surface of MNPs was investigated by Fourier-transform infrared (FT-IR) spectra on a Nicolet iS10 FT-IR spectrometer (Thermo Scientific^TM^, USA). MNPs were freeze-dried (FreeZone 4.5 L benchtop freeze dryer, Labconco Corporation, USA) and prepared as potassium bromide (KBr) discs. Corresponding spectra were collected in the range of 500 – 4000 cm^-1^. The background spectra were measured by the clean KBr discs containing no sample material.

**1.3 Fundus fluorescein angiography (FFA) and optical coherence tomography (OCT) imaging procedure**

To examine the morphological changes after laser photocoagulation, fluorescein angiography was carried out as described previously.^1, 2^ In brief, the mice were anesthetized and intraperitoneally injected with 1 % AK-FLUOR (Alcon, 100 cc 20 g^-1^ mouse) to obtain a clear bright field image. The mice were then placed on the platform of the imaging system of Micron III fundoscopy system (Phoenix Research Laboratories, Pleasanton, CA, USA) and the position was adjusted until a clear image of the fundus filled the screen and the OCT was visualized. The area of interest was acquired using StreamPix software.

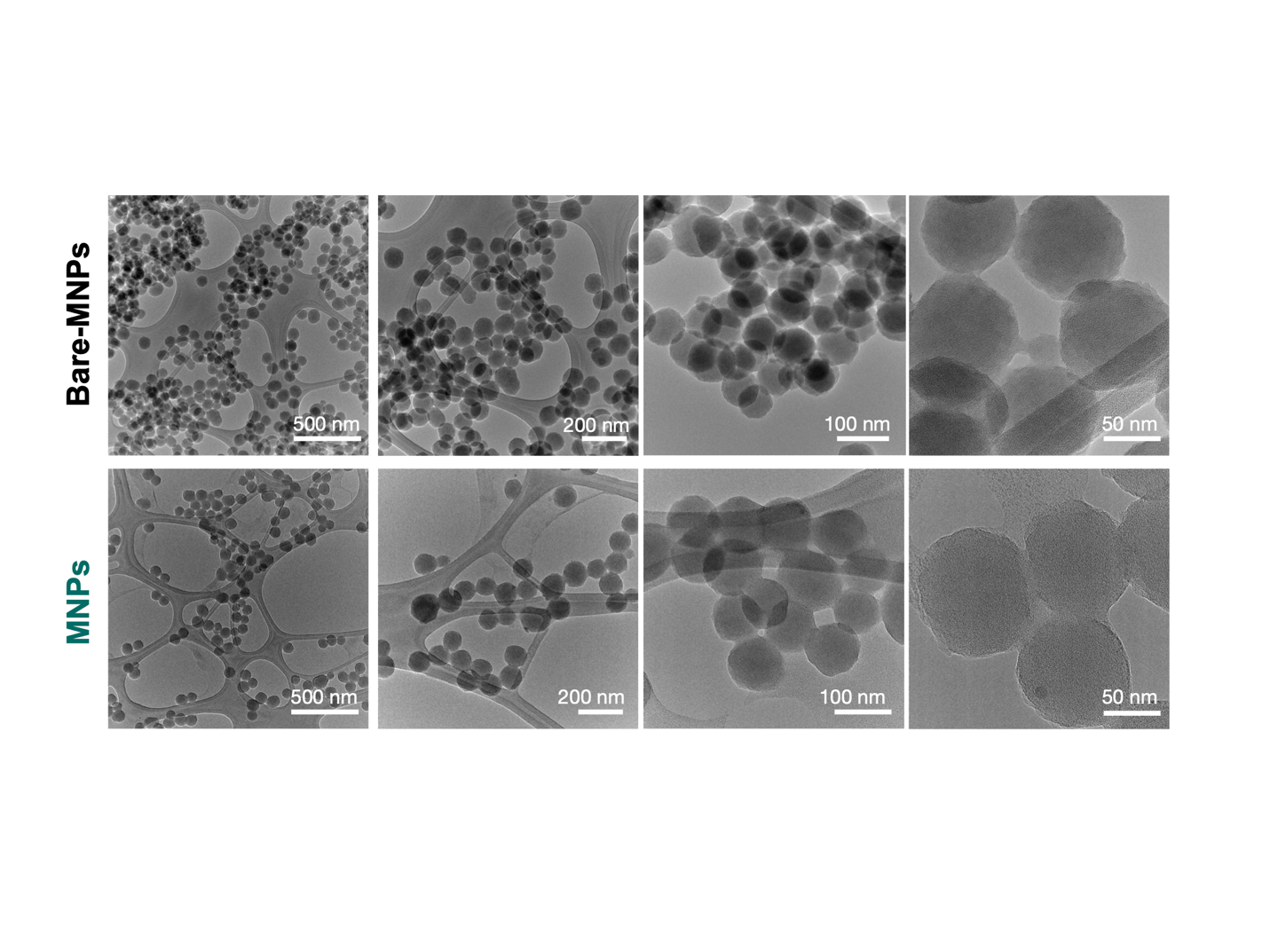

Figure S 1. TEM images of Bare-MNPs and MNPs at different magnifications. The shape and size of nanoparticles were analyzed by TEM. There were no significant changes to the nanoparticles after surface modification with mPEG-SH.

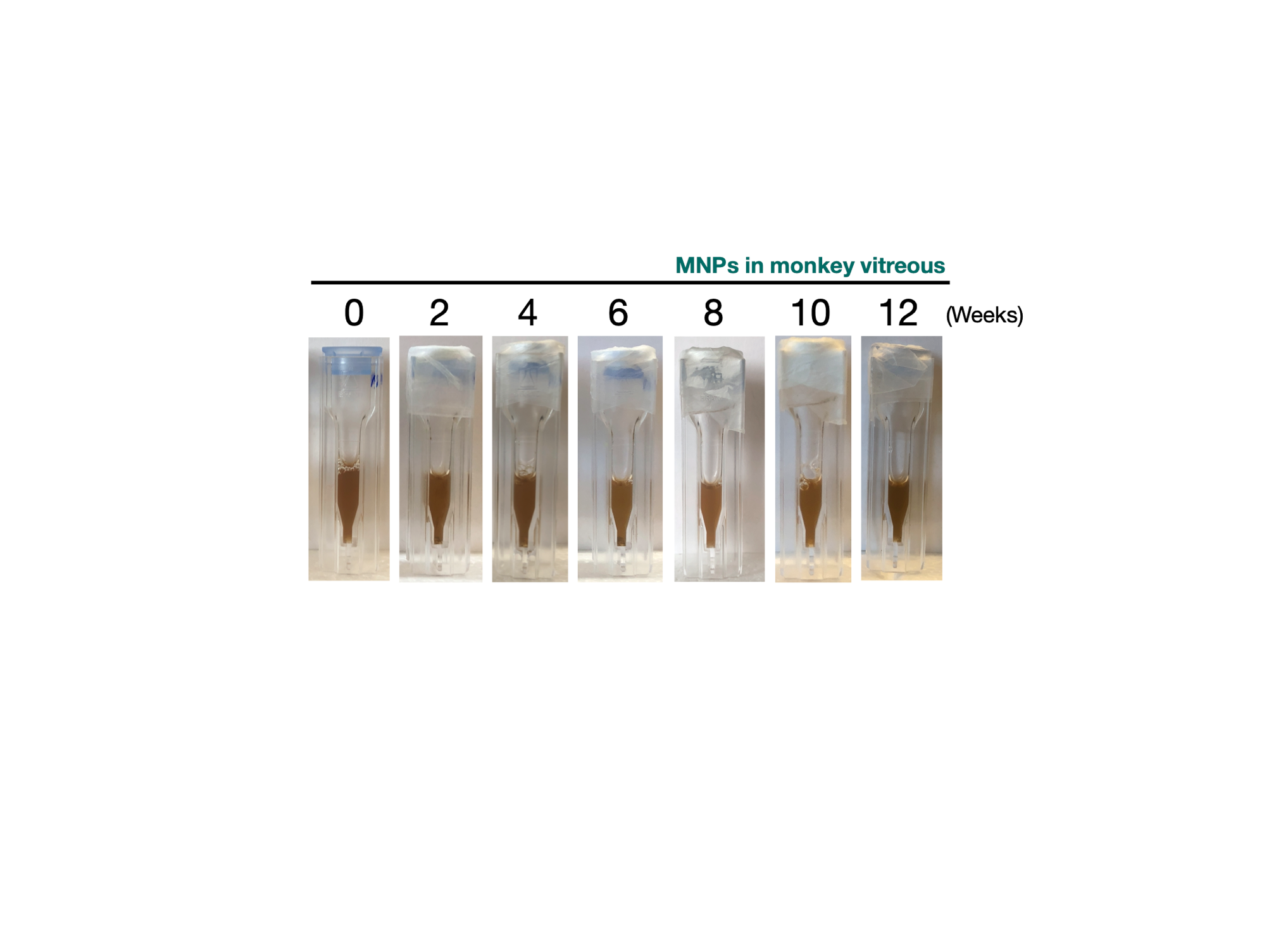

Figure S 2. Photographs of MNPs dispersed in monkey vitreous at different times. Results showed that the colloidal stability was maintained long-term (12 weeks) in monkey vitreous.

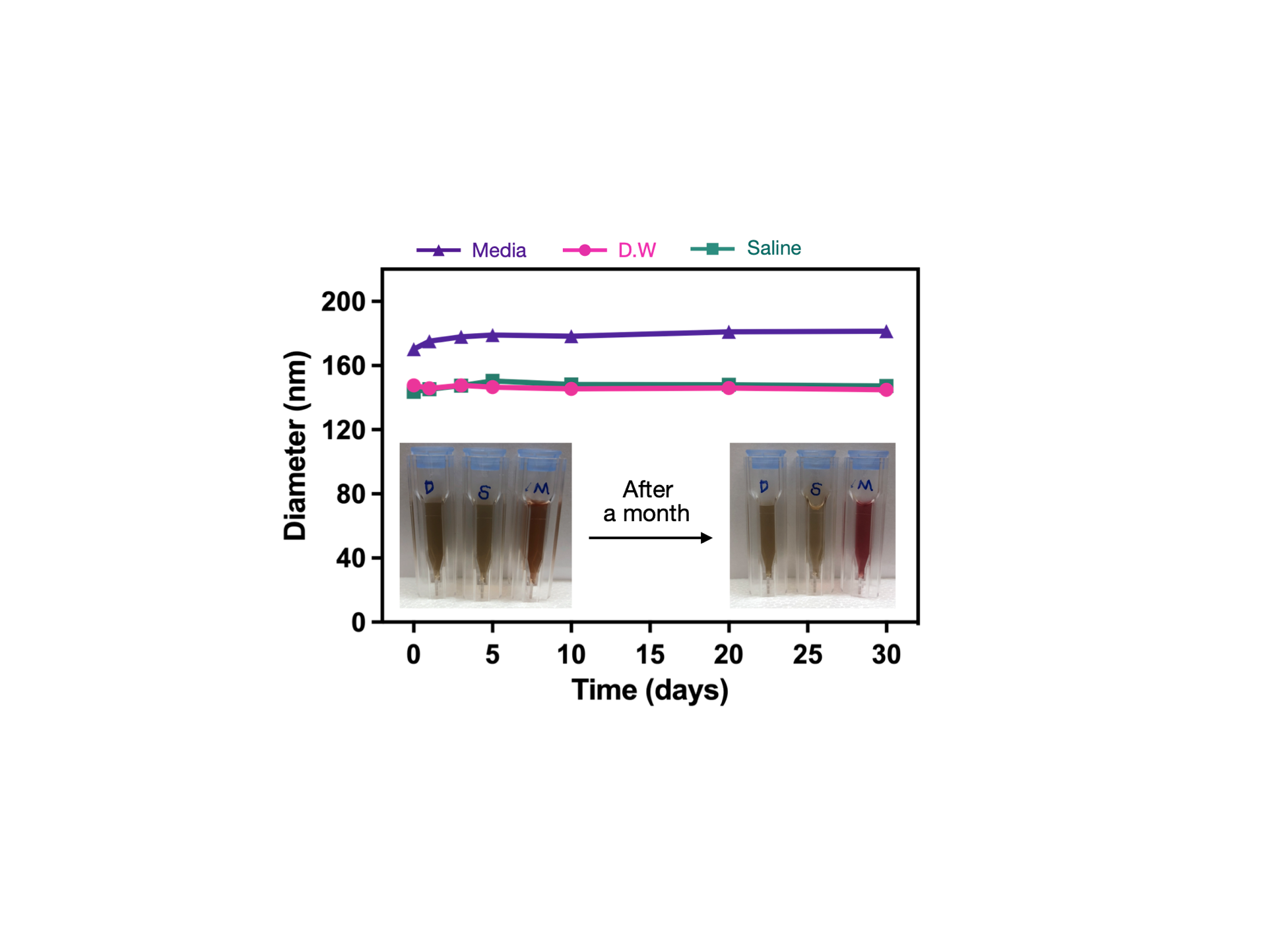

Figure S 3. MNPs show excellent colloidal stability in different solutions. The size of MNPs was determined by DLS. The inserted pictures showed MNPs in D.W, saline, and cell culture media at 0 vs 30 days.

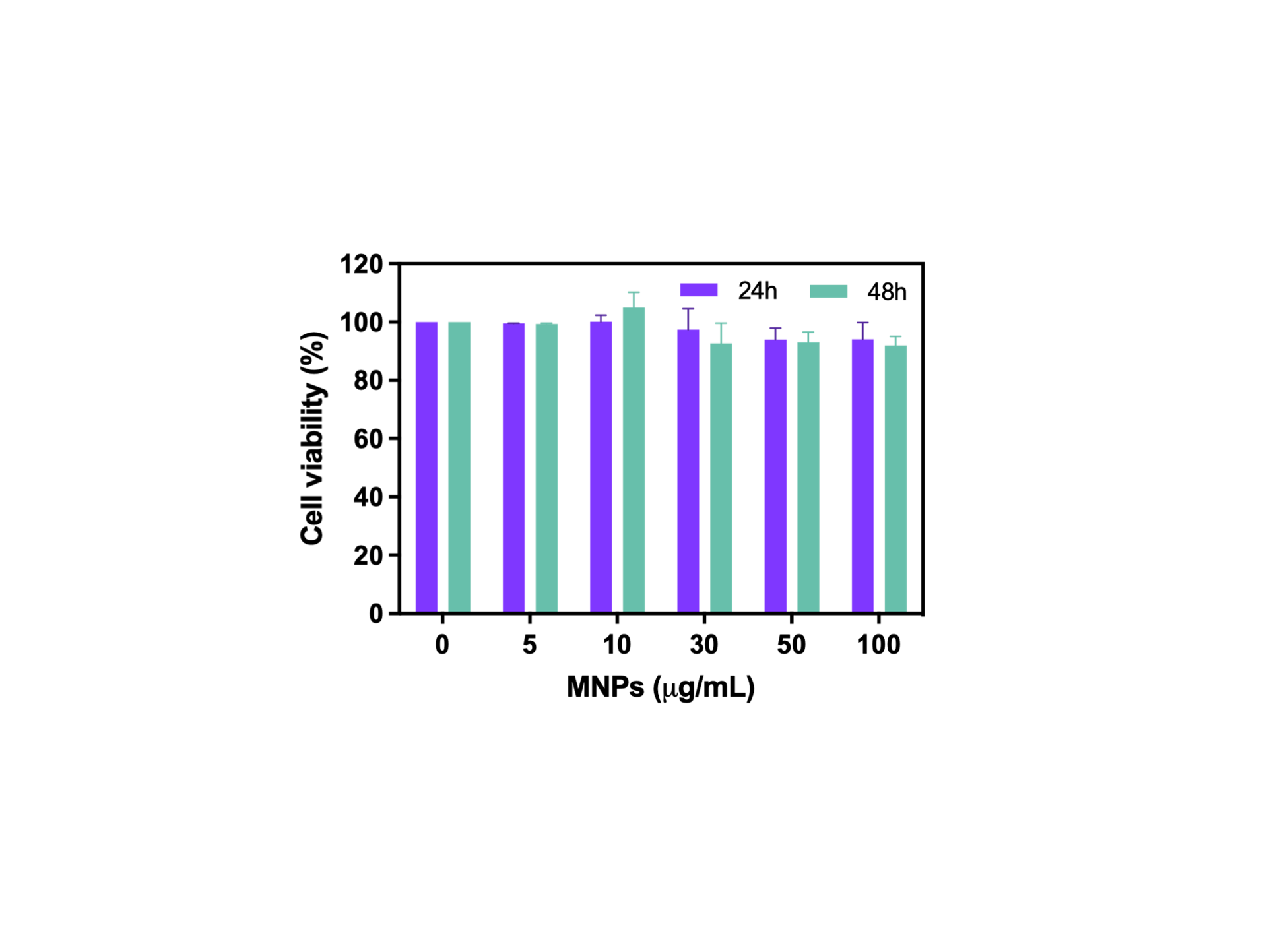

Figure S 4. Cell cytotoxicity assessment of MNP_S_ in ARPE19 cells. MNPs showed high cell viability (< 90 %) in ARPE19 cells after treatment with increasing concentration from 0 to 100 μg mL^-1^ for 24 and 48 h.

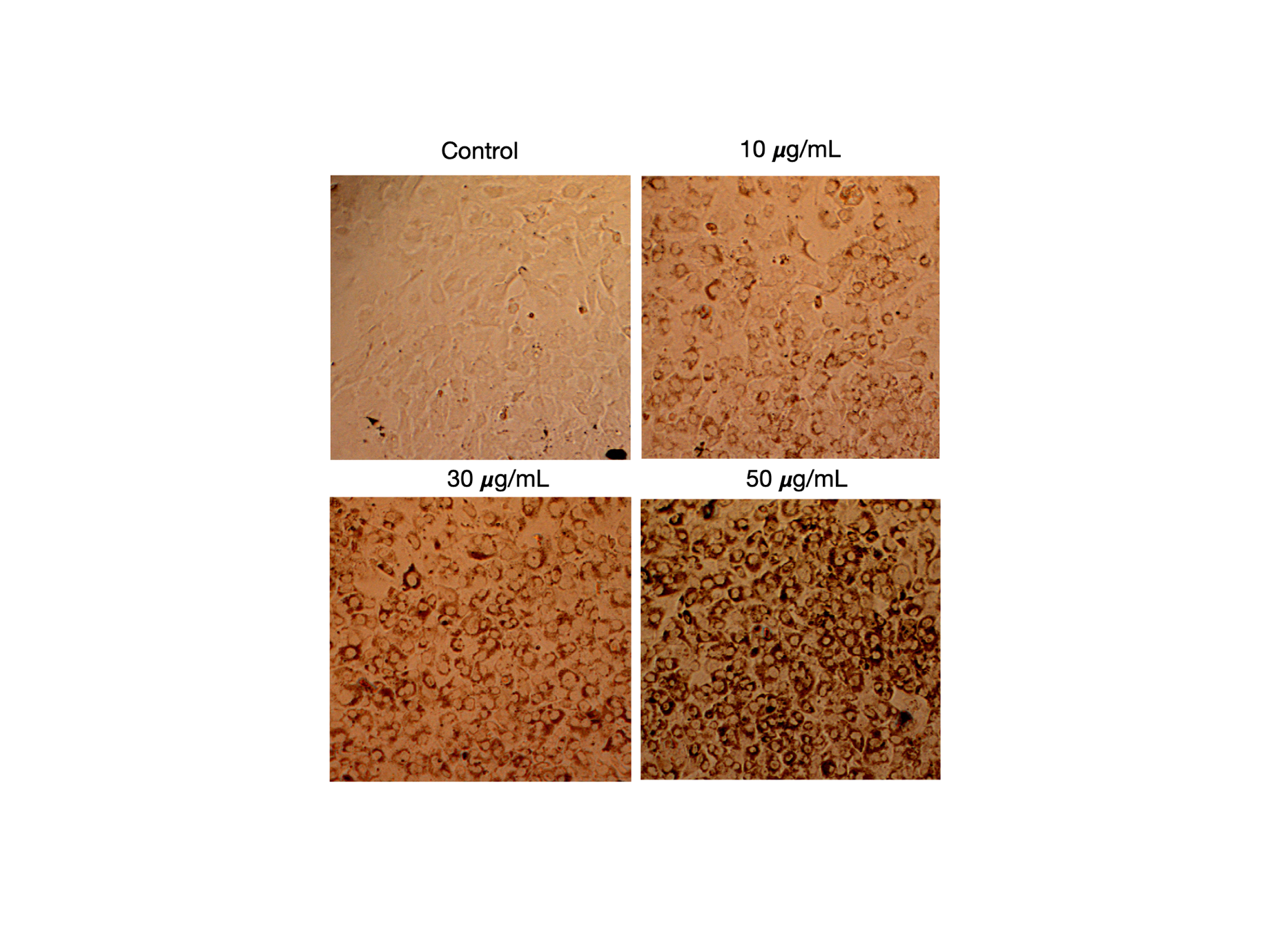

Figure S 5. Representative bright-field images (10x magnification) showing intracellular uptake of MNPs by ROS-induced ARPE19 cells after 12 h incubation.

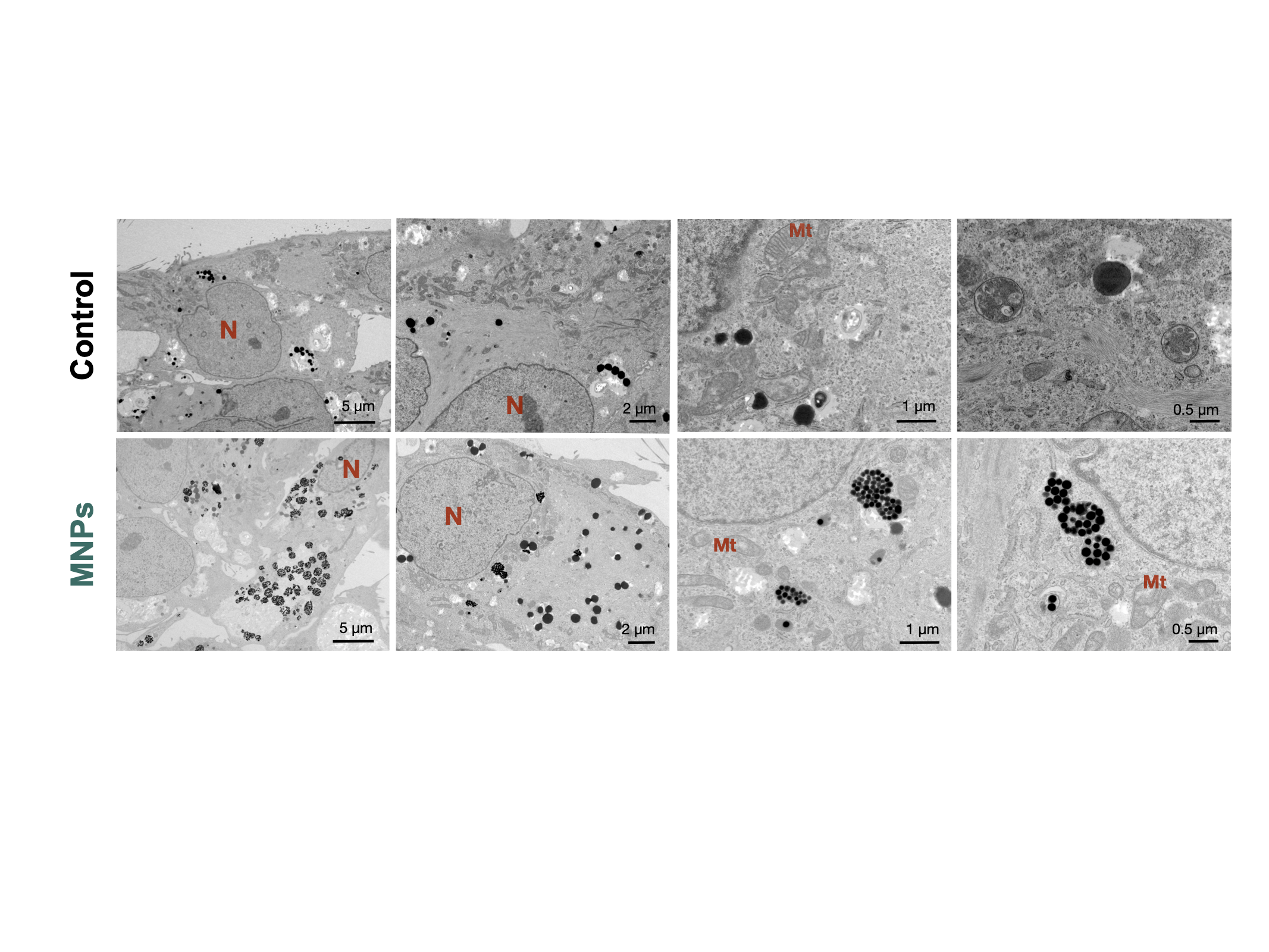

Figure S 6. Visualization of the intracellular uptake and trafficking of MNPs in ARPE19 cells. Bio-TEM images of ARPE19 cells incubated with MNPs for 24 h. All cells were fixed with 4 % formaldehyde and osmium tetroxide, then embedded in epoxy resin and stained with 4 % uranyl acetate (pH = ~5, 5 min of staining, rapid drying) to enhance the image contrast. N: nuclear, Mt: Mitochondria.

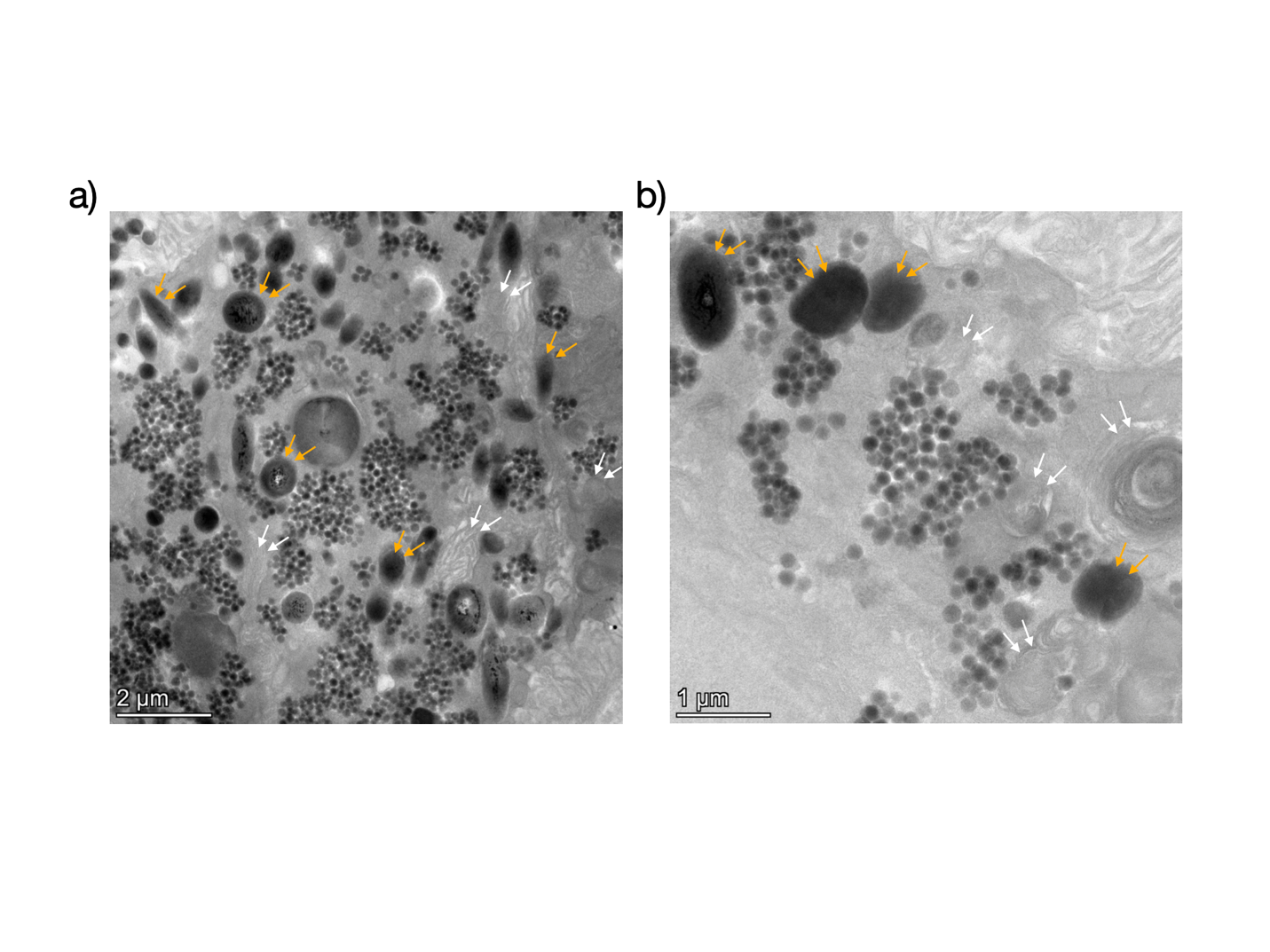

Figure S 7. Visualization of the intracellular uptake and trafficking of MNPs in the RPE layer at 15 days after single-dose IVT injection. The Bio-TEM images at a) low and b) high magnification show that MNPs are clearly visible in the cytosol of the RPE in MNPs-treated (1 μL, 10 μg μL^-1^) eyes, but not in the mitochondria and melanosomes. All ocular tissues were stained with 4 % uranyl acetate solution (pH = ~5, 5 min of staining, rapid drying) to enhance the image contrast. The white-colored and yellow-colored arrows indicate the mitochondria and melanosomes, respectively.
